# Triplet-Triplet Annihilation PLGA-Nanoparticles for Cancer Bioimaging

**DOI:** 10.1101/2020.09.02.274969

**Authors:** Olena Vepris, Christina Eich, Yansong Feng, Hong Zhang, Eric. L. Kaijzel, Luis J. Cruz

**Affiliations:** Department of Radiology, Leiden University Medical Center (LUMC); Van ‘t Hoff Institute for Molecular Sciences. University of Amsterdam, Science Park 904, 1098 XH, Amsterdam, the Netherlands

**Keywords:** Photon upconversion, Triplet-triplet annihilation, *In vivo* imaging, PLGA, nanoparticles

## Abstract

Triplet-triplet annihilation upconversion (TTA-UC) nanoparticles (NPs) have emerged as imaging probes and therapeutic probes in recent years due to their excellent optical properties. In contrast to lanthanide ions-doped inorganic materials, highly efficient TTA-UC can be generated by low excitation power density, which makes it suitable for clinical applications. In the present study, we used biodegradable poly(lactic-co-glycolic acid) (PLGA)-NPs as delivery vehicle for TTA-UC based on the heavy metal porphyrin Platinum(II) octaethylporphyrin (PtOEP) and the polycyclic aromatic hydrocarbon 9,10-diphenylanthracene (DPA) as photosensitizer/emitter pair. TTA-UC-PLGA-NPs were successfully synthesized according to an oil-in-water emulsion and solvent evaporation method. After physicochemical characterization, UC-efficacy of TTA-UC-PLGA-NPs was assessed *in vitro* and *ex vivo*. TTA-UC could be detected in the tumor area 96 hours after *in vivo* administration of TTA-UC-PLGA-NPs, confirming the integrity and suitability of PLGA-NPs as TTA-UC *in vivo* delivery system. Thus, this study provides proof-of-concept that the advantageous properties of PLGA can be combined with the unique optical properties of TTA-UC for the development of advanced nanocarriers for simultaneous *in vivo* molecular imaging and drug delivery.

## Introduction

Since the introduction of imaging techniques to diagnosis and treatment of different pathologies, great progress has been achieved in the field of medical imaging, and the variety of imaging agents become more sophisticated in terms of efficacy, safety and target-specificity ^1^.

In the last decade, photon upconversion (UC), a process that converts low-energy into high-energy light ^2,3^, has become state-of-art in the field of *in vitro* and *in vivo* biomedical imaging due to several unique optical properties: i) large hypsochromic shift, ii) sharp emission peak, iii) long luminescence shelf-life and iv) high photostability ^4-6^. Photon UC is a combination of photophysical processes and it can be achieved via different energy transfer mechanisms using different materials, such as rare-earth metals, metalloporphyrins and organic polyaromatic hydrocarbons ^2,3^.

Triplet-triplet annihilation (TTA) is a special form of UC, resulting in superior optical properties, such as intense absorption of excitation light and high UC quantum yield, while using low excitation power density ^3,7,8^; the latter makes TTA-UC in particular interesting for biomedical applications. In TTA-UC, the energy transfer between a sensitizer and an emitter molecule takes place through a series of subsequent nonradiative mechanisms, such as intersystem crossing (ISC) and triplet-triplet energy transfer (TTET) (Figure 1)^3,7,8^.

**Figure 1.**
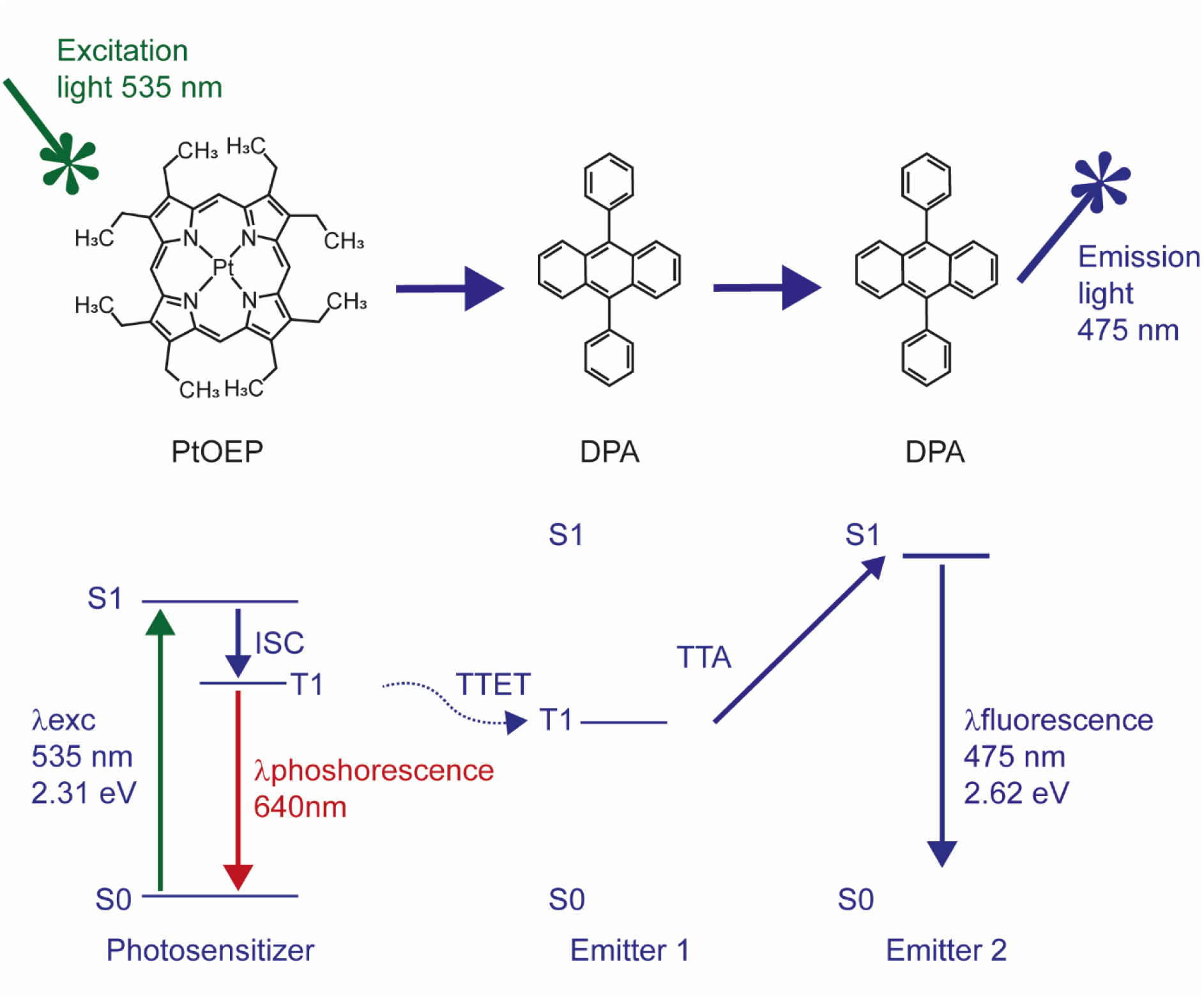
Jablonski diagram describing the energy levels of the electrons involved in energy transfer between PtOEP and DPA molecules. The energy values were calculated based on theoretical data.

When electromagnetic radiation “hits” a sensitizer molecule its electrons will be promoted from the ground to a singlet excited state, and then transferred to a triplet excited state nonradiatively. The released energy that accompanies the transition of the triplet excited electron to the ground singlet state will be transferred to an emitter/acceptor molecule to pump the latter to an excited triplet state. If two of these triplet excited state molecules are close enough, the interaction between the two will cause the transition of one of them to the ground singlet state and the other to a singlet excited state, *i*.*e*. the collision results in an annihilation of the triplet excited state of one acceptor molecule and promotion of another one to the singlet excited state (Figure 1).

The UC-process is a highly improbable event according to the quantum mechanical principle, since it consists of a series of spin-forbidden transitions ^9^. It means that the UC-process should be boosted under favourable conditions that promote those transitions. For instance, to promote ISC, which is the transition of a molecule from a singlet electronic excited state to a triplet electron state, it is advantageous to choose molecules with strong spin-orbit coupling and an expanded π conjugated system. Metalloporphyrins, such as the green absorbing platinum octaethylporphyrin (PtOEP), possess both requirements due to the presence of a transition metal that promotes a spin-forbidden transition and a cyclic π conjugated system that enhances ISC yield ^10,11,12^. More recently, lanthanide complexes of porphyrinoids inside of nanomicelles and mesoporous silica NPs were employed for in vivo imaging in HeLA cells, demonstrating the potential of porphyrinoids as sensitizers in TTA-UC ^13^. In the present study, we describe an approach whereby TTA-UC is combined with biocompatible poly(lactic-co-glycolic acid) (PLGA) to create an UC-PLGA-nanoparticle (NP) system as a potential tool for cancer visualization. NPs, such as those made of biodegradable and FDA-approved PLGA, protect their payload from premature degradation ^14-20^, are well described, chemically adaptable and can guide their payload inside target cells *in vitro* and *in vivo* ^19-22^.

To generate TTA-UC, we selected PtOEP as the photosensitizer and 9,10-diphenylanthracene (DPA) as the annihilator; a combination that has been reported to result in highly efficient UC yield ^23,24^. PtOEP and DPA were successfully encapsulated into the hydrophobic core of PLGA-NPs. Phosphorescence and UC fluorescence signals resulting from TTA-UC-PLGA-NPs could be detected *in vitro, in vivo* and *ex vivo* up to 96 hours after injection. We conclude that PLGA provides a suitable environment for TTA-UC, which could be utilized for simultaneous optical imaging and drug delivery *in vivo*.

## Results and Discussion

### Preparation, physicochemical characterization and cytotoxicity of TTA-UC-PLGA-NPs

In the present study, TTA-UC-PLGA-NPs were successfully synthesized applying an oil-in-water emulsion and solvent evaporation method, as previously described ^19,20,22,25^. Briefly, 100 mg of PLGA and 1 mg of each chromophore were dissolved in 3 ml of dichloromethane (DCM). The mixture of chromophores and PLGA polymer was emulsified under sonication in the presence of PVA. After removal of the organic solvent, the TTA-UC-PLGA-NPs were collected by centrifugation, washed and freeze-dried. The oil-in-water emulsion and solvent evaporation method led to an encapsulation efficiency (EE) of PtOEP and DPA molecules of 24% and 40%, respectively, as determined by nanodrop measurement of dissolved NPs (data not shown).

Due to the high sensitivity of the porphyrin triplet states to oxygen quenching, the actual lifetime of the triplet state depends on the solvent, its purity and on its oxygen content ^26^. To protect the optical properties of chromophores, PtOEP and DPA were dissolved in DCM. To ensure that encapsulation into PLGA-NPs did not affect the optical properties of PtOEP and DPA, TTA-UC-PLGA-NPs were dissolved in dimethylformamide (DMF) and analysed by absorption curve analysis and compared to pure compounds (Figure 2). The absorption spectrum of PtOEP depicts the characteristic shape of porphyrin absorption: a strong absorption peak at 380 nm (Soret band) was accompanied by a weaker absorption peak located at 501 and 533 nm (Q-band) ^27^ (Figure 2A). DPA, the emitter for the TTA-UC system, showed several absorption peaks between 350 nm and 400 nm (Figure 2B). When the TTA-UC-PLGA-NPs were dissolved in DMF, the absorption spectrum showed the combined characteristic optical profiles of PtOEP and DPA (Figure 2C). In conclusion, absorption spectrum analysis of TTA-UC-PLGA-NPs confirmed the successful encapsulation of both chromophores into the PLGA polymer, while the optical properties of PtOEP and DPA were maintained.

**Figure 2.**
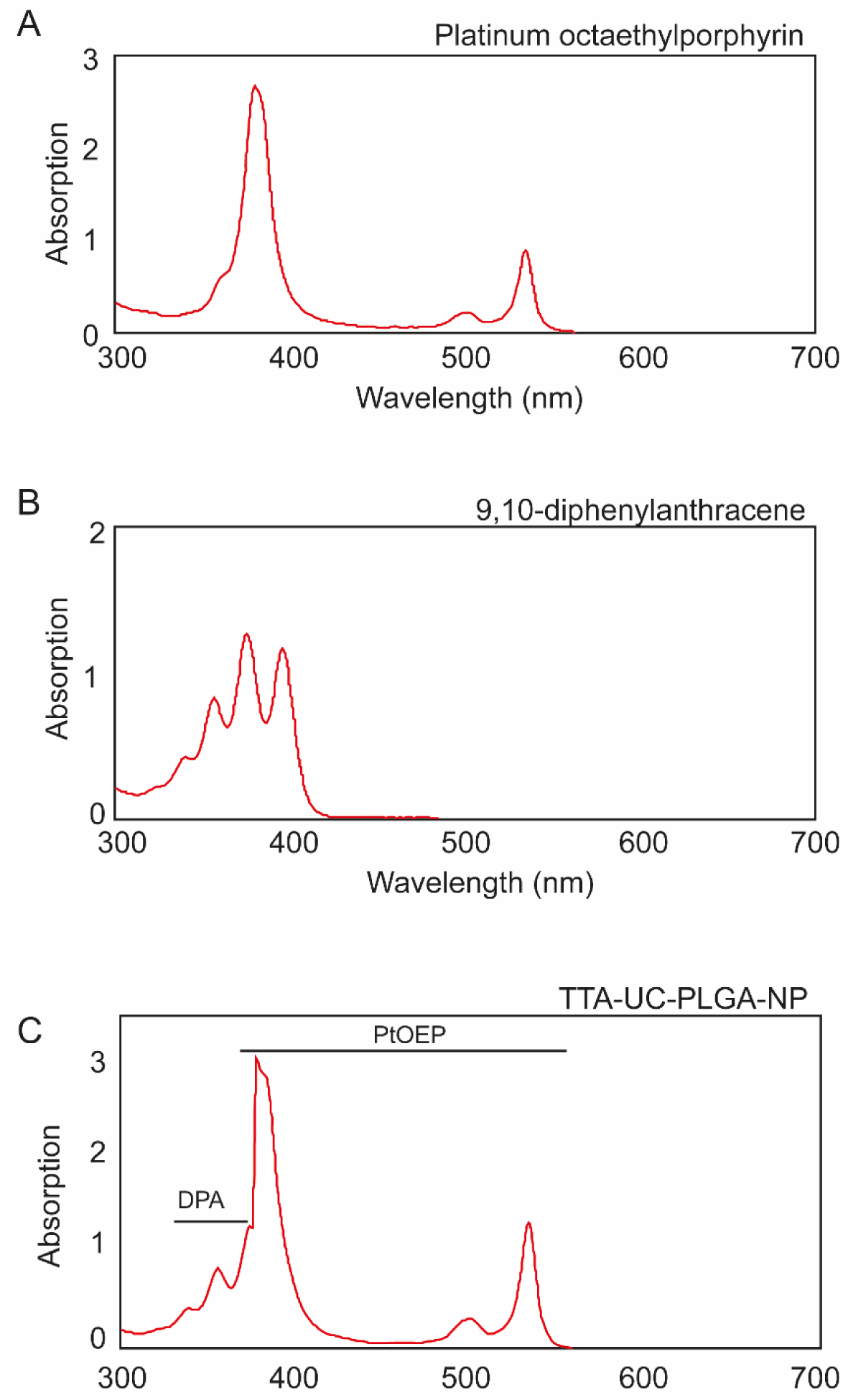
Individual absorption spectra of (A) platinum octaethylporphyrin (PtOEP) and (B) 9,8-diphenylanthracene (DPA) each dissolved in dimethylformamide (DMF). (C) TTA-UC-PLGA-NPs were hydrolysed overnight in 0.8M NaOH at 37°C and the absorption spectrum of co-encapsulated PtOEP and DPA were measured.

The size of the NPs was determined by dynamic light scattering (DLS) analysis as ∼200 nm in diameter (Table 1, Figure 3A) and the polydispersity index (PDI) value of 0.4 obtained from the DLS measurement indicated a homogeneous size distribution (Table 1).

**Table 1.** Physicochemical properties of NPs. Data are present as average ± SD (N=3)

| NPs | Size (nm) | Zeta potential (mV) | PDI | Loading (µg/mg) | Encapsulation efficiency (%) |
| --- | --- | --- | --- | --- | --- |
| TTA-UC PLGA | 197.8 ± 47.5 | -30.7 ± 5.0 | 0.4 | PtOEP: 244.0<br>DPA: 396.0 | PtOEP: 24.4<br>DPA: 39.6 |

**Figure 3.**
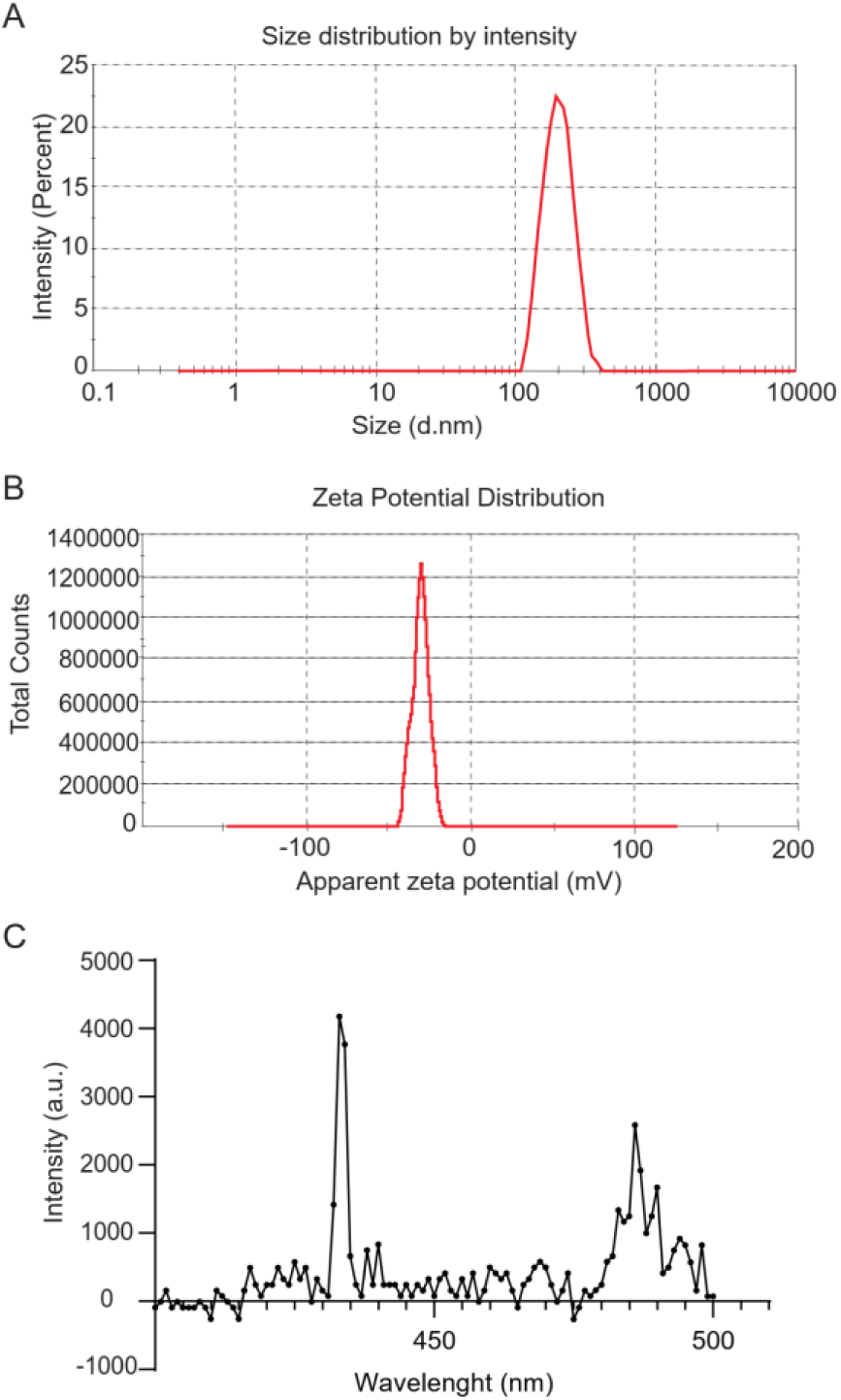
Physicochemical characterization of TTA-UC-PLGA-NPs. Representative (A) dynamic light scattering and (B) zeta-potential measurement of TTA-UC-PLGA-NPs reconstituted in water after freeze-drying. The average size of the nanoparticles was 198 nm in diameter and the average zeta potential value was −31mV. (C) TTA-UC-PLGA-NPs were dissolved in water and directly analysed. The NPs in solution were excited at 535 nm and the emission was collected using a photofluorometer.

NPs smaller than 10 nm are known to be eliminated by renal excretion ^28^, while NPs with sizes ranging from 50 to 300 nm showed extended circulation times in the blood stream compared to NPs of larger size ^29^. Hence, TTA-UC-PLGA-NPs of 200 nm have the ideal size for prolonged systemic circulation times; an important prerequisite to reach the tumor site. In addition, ZetaSizer measurement showed that the surface charge of the NPs was on average −30 mV (Table 1, Figure 3B). The surface charge is a crucial factor to determine the stability of the NPs in solution, but also affects cellular uptake, biodistribution and cytotoxicity. A high surface charge guarantees an electrostatic stabilization of the NPs due to a strong surface repulsion between NPs of the same charge. According to the measured value, it is possible to assume that the NPs are stable in suspension for a prolonged period of time _30,31_.

Next, we investigated the optical properties of the TTA-UC system after encapsulation into PLGA-NPs. To this purpose, TTA-UC-PLGA-NPs were dissolved in water and excited at 535nm (Figure 3C). The TTA-UC signal could be detected at 433 nm, confirming the integrity of the TTA-UC system after encapsulation into PLGA.

The charge of UC-NPs has been shown to affect the intracellular localization and cellular cytotoxicity ^32^. Positively charged UC-NPs localized to mitochondria, while negatively charged UC-NPs preferentially localized to lysosomes and the cytoplasm, which was associated with lower cellular cytotoxicity ^32^. To assess potential toxicity of our NPs, TTA-UC-PLGA-NPs were incubated at different concentrations (25-200 µg/ml) for 72 hours with OVCAR-3 ovarian cancer cells, and the cellular toxicity was determined by MTS assay (Figure 4). TTA-UC-PLGA-NPs were compared to empty control PLGA-NPs. No toxicity of TTA-UC-PLGA-NPs or empty PLGA-NPs on OVCAR-3 cells was observed after 72 hours (Figure 4).

**Figure 4.**
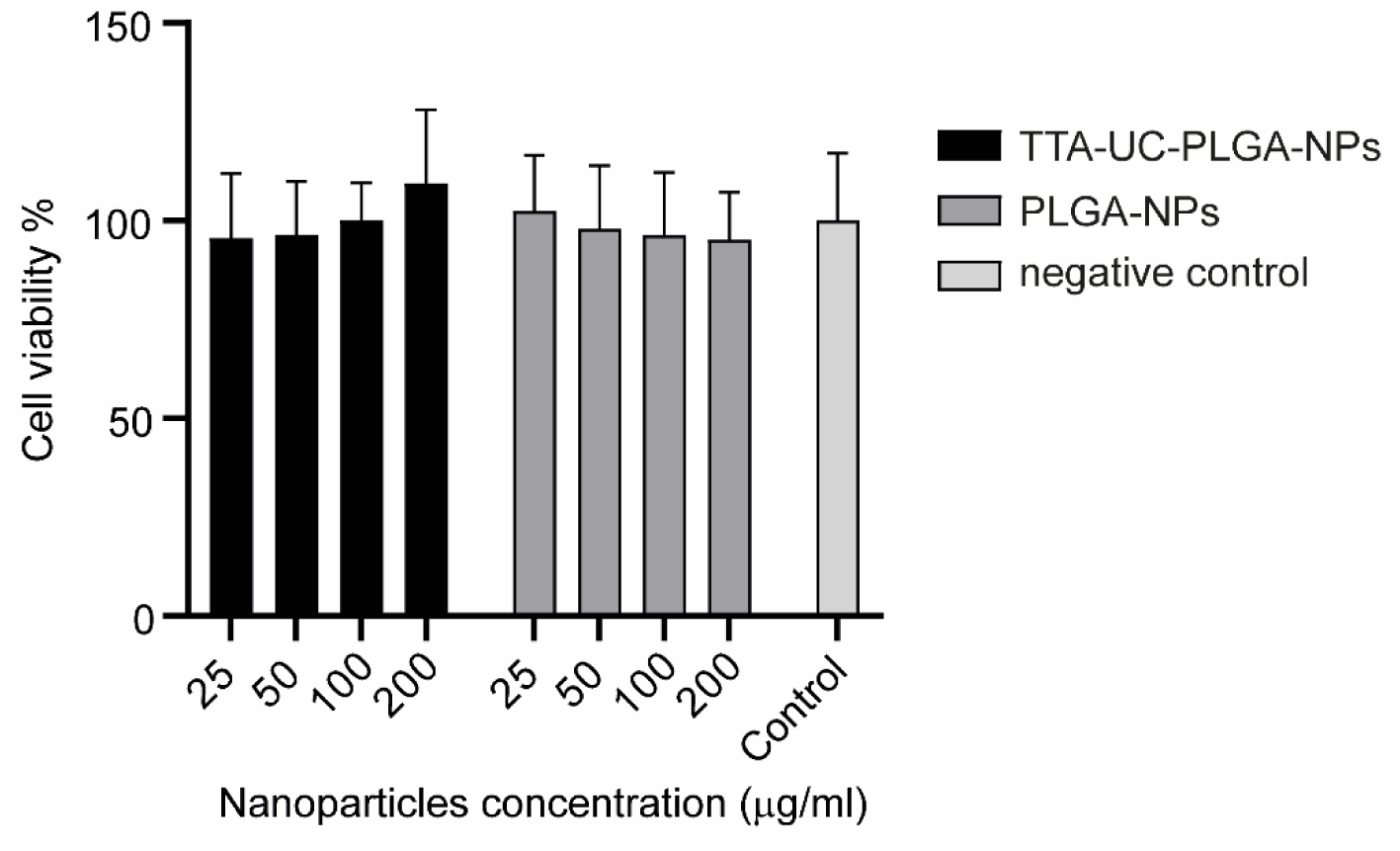
Assessment of cellular cytotoxicity of TTA-UC-PLGA-NPs. OVCAR-3 cells were incubated with 25, 50, 100 and 200 µg/ml TTA-UC-PLGA-NPs or control PLGA-NPs for 72 hours. Untreated cells were used as negative control. The cell viability was assessed after 72 hours by MTS assay.

### Cellular Uptake of TTA-UC-PLGA-NPs characterized by fluorescence microscopy

To assess the cellular uptake of TTA-UC-PLGA-NPs, OVCAR-3 cells were incubated for 2 hours with 200 μg/ml TTA-UC-PLGA-NPs and analysed by fluorescent microscopy. In order to confirm the intracellular localization of the NPs, the cells were counterstained with DAPI (λexcitation = 340-380 nm, λemission = 425 nm) and DiD (λexcitation = 676-688 nm, λemission = 700-742 nm) for nucleus and membrane detection, respectively. To detect the TTA-UC-PLGA-NPs, the samples were excited at 542-582 nm, and the phosphorescence signal was collected at 604-644 nm (Figure 5). TTA-UC-PLGA-NPs were successfully taken up by OVCAR-3 cells. The overlay of the membrane staining with the phosphorescence signal of the TTA-UC-PLGA-NPs confirmed the intracellular localization of the NPs (Figure 5).

**Figure 5.**
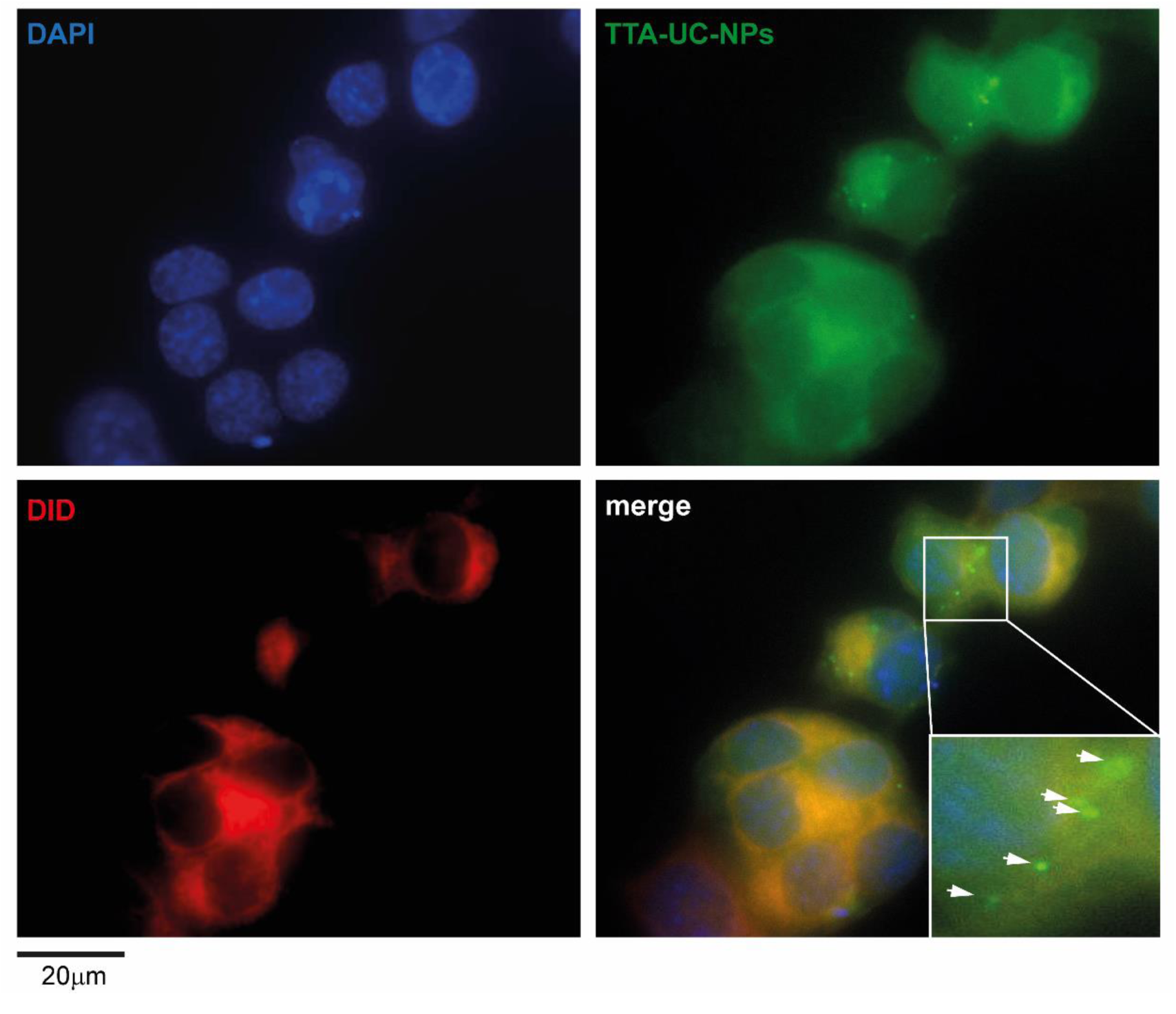
Cellular uptake of TTA-UC-PLGA-NPs. OVCAR-3 cells were incubated with 200 µg/ml TTA-UC-PLGA-NPs for 2 hours at 37°C and analysed by fluorescence microscopy. The cells were fixed, and the nucleus was stained with DAPI (blue) and the cell membrane was stained with DiD (red). The phosphorescence signal (green) of the TTA-UC-PLGA-NPs was collected. The following filter cubes were used to discriminate DAPI, DiD and TTA-UC-PLGA-NP-signals: DAPI, λexc = 340-380 nm and λemission = 425nm); DiD (λexc = 676-688 nm, λemission = 700-742nm; TTA-UC-PLGA-NPs, λexc = 542-582 nm, λemission 604-644 nm. Scale bar = 20µm.

### Characterization of TTA-UC-PLGA-NPs UC properties

The ability of TTA-UC-PLGA-NPs to generate the UC process was assessed by confocal microscopy (Figure 6). The NPs were excited at a wavelength of 535 nm, and the emission signals were collected between 430-475 nm and 650-670 nm. In our study, the emission shift towards shorter wavelengths (hypsochromic shift) went from green (535 nm) to blue (430-475 nm) (Figure 6). It means that the energy associated with the emission wavelength was higher than the energy associated with the excitation wavelength, as a consequence of the anti-Stokes shift in the TTA-UC process ^33^. On the contrary, the change in emission spectrum towards longer wavelengths (bathochromic shift), in our case 650-670 nm, is a characteristic of all conventional luminescence processes (Figure 6). Using confocal microscopy, we could image single TTA-UC-PLGA-NPs that created dual emission wavelengths: the red emission from the phosphorescence of PtOEP and the blue emission as a result of the TTA-UC process. Thus, we confirmed the preparation of functional TTA-UC-PLGA-NPs.

**Figure 6.**
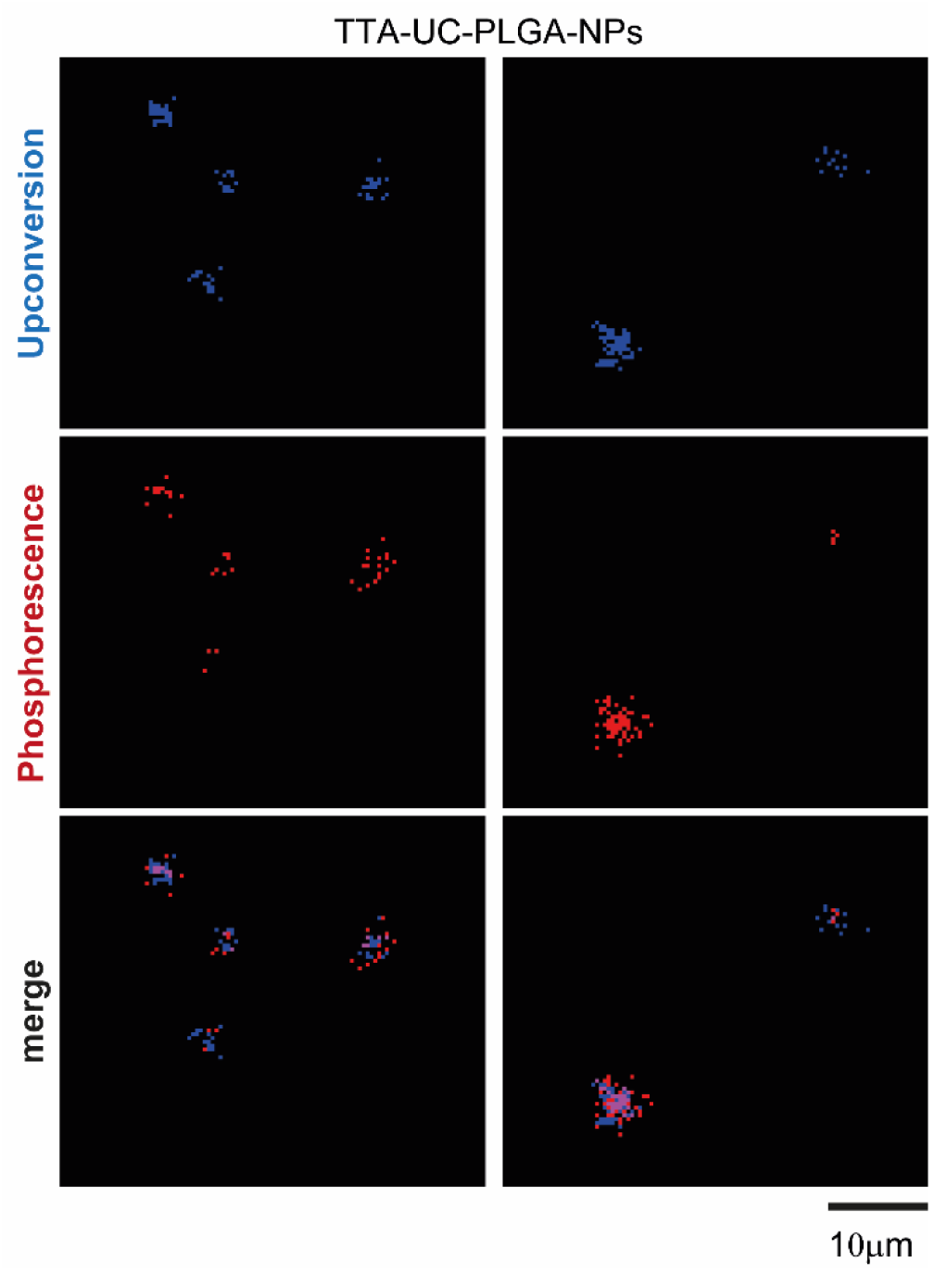
TTA-UC imaged by confocal microscopy. UC-TTA-PLGA-NPs were dissolved in water and imaged on a glass cover slip. The NPs were excited at 535 nm and the following emission wavelengths were collected: 430-475 nm (blue) = upconversion, 650-670 nm (red) = phosphorescence. Scale bar = 10µm.

The dual colour feature of TTA-UC-PLGA-NPs was useful to improve signal-to-noise ratio and to estimate the ratio of UC/phosphorescence signal. The overall efficiency of the TTA-UC process depends on each elementary step (ISC, TTET, TTA), on the concentration of emitter molecules and the presence of other competing molecules, such as elementary oxygen ^7,26^. Additional challenges are encountered with when TTA-UC is integrated in a NP-system. Aggregation of photosensitizers and emitters can have detrimental effects on the optical properties of TTA-UC-NPs and, as a consequence, the loss of mobility of photosensitizers/emitters can limit the UC-process ^7,34^. Thus, creating a highly efficient UC-NP-system without any secondary decay remains an enormous challenge for practical application ^35^.

### Stability assessment of TTA-UC-PLGA-NPs in solution

As PLGA-polymers are subject to hydrolysis in a water environment ^36^ we investigated the stability of the optical properties of TTA-UC-PLGA-NPs over time (Figure 7). To this purpose, TTA-UC-PLGA-NPs were dissolved in PBS and incubated at 37°C in shaking mode. At time point 0, after 24 and 72 hours, the dispersion of TTA-UC-PLGA-NPs was excited at 535 nm and the TTA-UC and phosphorescence processes were measured at 433 and at 642 nm, respectively (Figure 7). The strongest phosphorescence and TTA-UC signals were detected at time point 0 (0 hours post dissolution) (Figure 7A-B). After 24 hours, the phosphorescence and TTA-UC signals decreased, but could still be detected after 72 hours incubation at 37°C. Thus, the data demonstrates that TTA-UC-PLGA-NPs are suitable for imaging processes lasting for at least 72 hours. The initial decrease in TTA-UC/phosphorescence intensity within the first 24 hours is likely the result of an early hydrolysis of the PLGA polymer, resulting in the partial leaking of the chromophores out of the PLGA-core. Interestingly, while the phosphorescence signal decreased 19-fold in the first 24 hours (determined at 640 nm), the TTA-UC signal only decreased 2-fold after 24 hours and 2.1-fold after 72 hours (determined at 433 nm). (Figure 7A-B). While both TTA-UC and phosphorescence intensities decreased over time, the TTA-UC signal relative to the phosphorescence signal increased, indicating that the TTA-UC process became more efficient. This might indicate that the initial hydrolysis of the PLGA polymer created a favourable environment for TTA-UC. The decrease in concentration of triplet excited emitter molecules due to degradation might have been counterbalanced by an increase in photosensitizer/emitter diffusion. Diffusion of sensitizer and emitter molecules is crucial to increase the probability of intermolecular TTET and TTA to take place ^26,34^.

**Figure 7.**
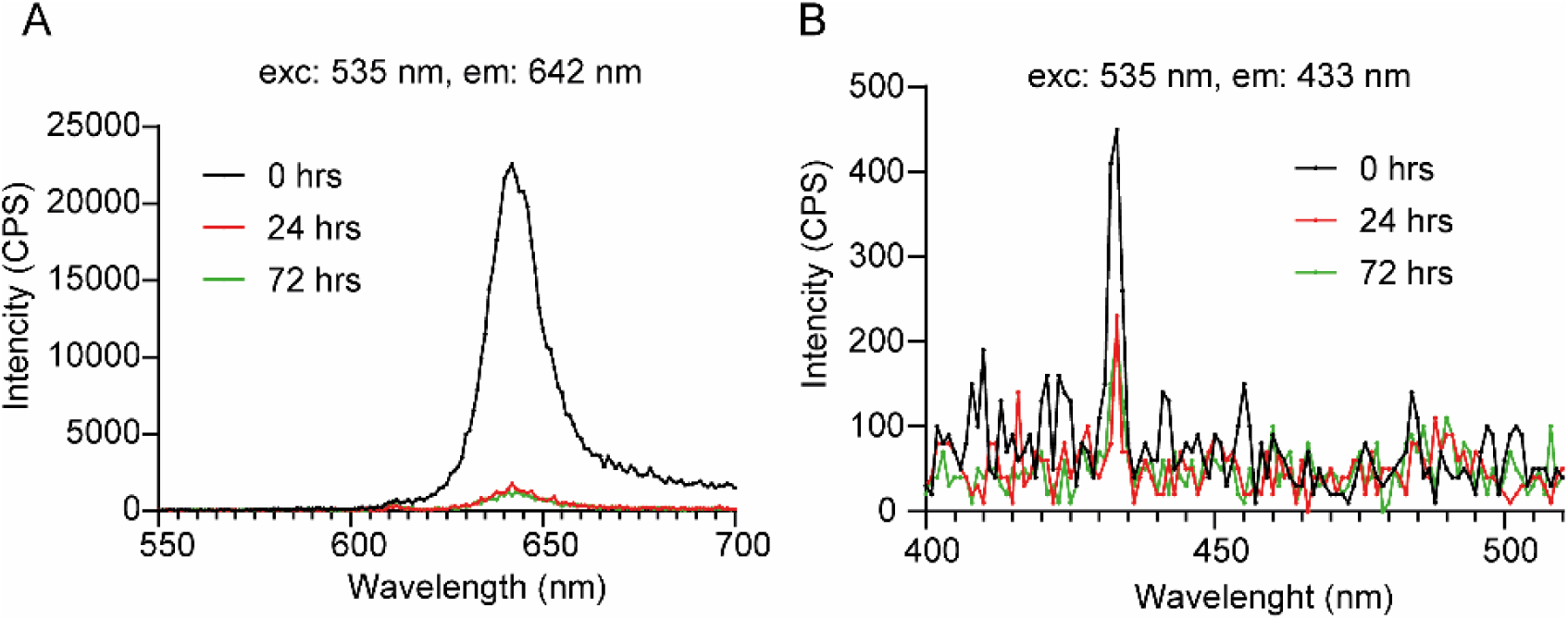
Stability assessment of TTA-UC-PLGA-NPs in solution. TTA-UC-PLGA-NPs were dissolved in water and incubated at 37°C for 72 hours. At time point 0, 24 and 72 hours, samples were analysed. The NPs in solution were excited at 535 nm and the emission was collected using a photofluorometer. (A) Phosphorescence and (B) TTA-UC emission graphs of TTA-UC-PLGA-NPs at various time points.

### In vivo monitoring of TTA-UC-PLGA-NPs

Real-time imaging of FVB mice was carried out with the goal to trace TTA-UC-PLGA-NPs in cancer cells *in vivo*. To this end, the IVIS imaging system was employed, which allowed measurement of the bathochromic emission captured at 640 nm (but not the UC process).

FVB mice were inoculated in the back with 1×10e^6^ human breast adenocarcinoma cells (MCF). When the tumors reached a volume of approximately 125 mm^3^, TTA-UC-PLGA-NPs were injected intravenously in the tail vein. At different time points post injection (3, 24, 48, 72 and 96 hours), the mice were imaged. *In vivo* imaging data showed that already 3 hours post injection, TTA-UC-PLGA-NPs accumulated in the tumor area (Figure 8A). The mice were monitored up to 96 hours post injection and the phosphorescence signal persisted during all imaged time points (Figure 8A-B). Quantification of the signal showed a significant increase in phosphorescence signal in the tumor area in the first 48 hours (p 0.026) and after 96 hours (p 0.0022), indicating that more TTA-UC-PLGA-NPs accumulated at the tumor site over time. Moreover, the data showed that once accumulated in the tumor, the NPs remained at the tumor site. Since the NPs did not present any specific targeting moiety for breast adenocarcinoma cancer cells, TTA-UC-PLGA-NPs most likely accumulated at the tumor site via the enhanced permeability and retention (EPR) effect. Tumor blood vessels are naturally leaky, with large openings up to 1.5 mm, thereby promoting the passive accumulation of NPs in tumors ^37 38^.

**Figure 8.**
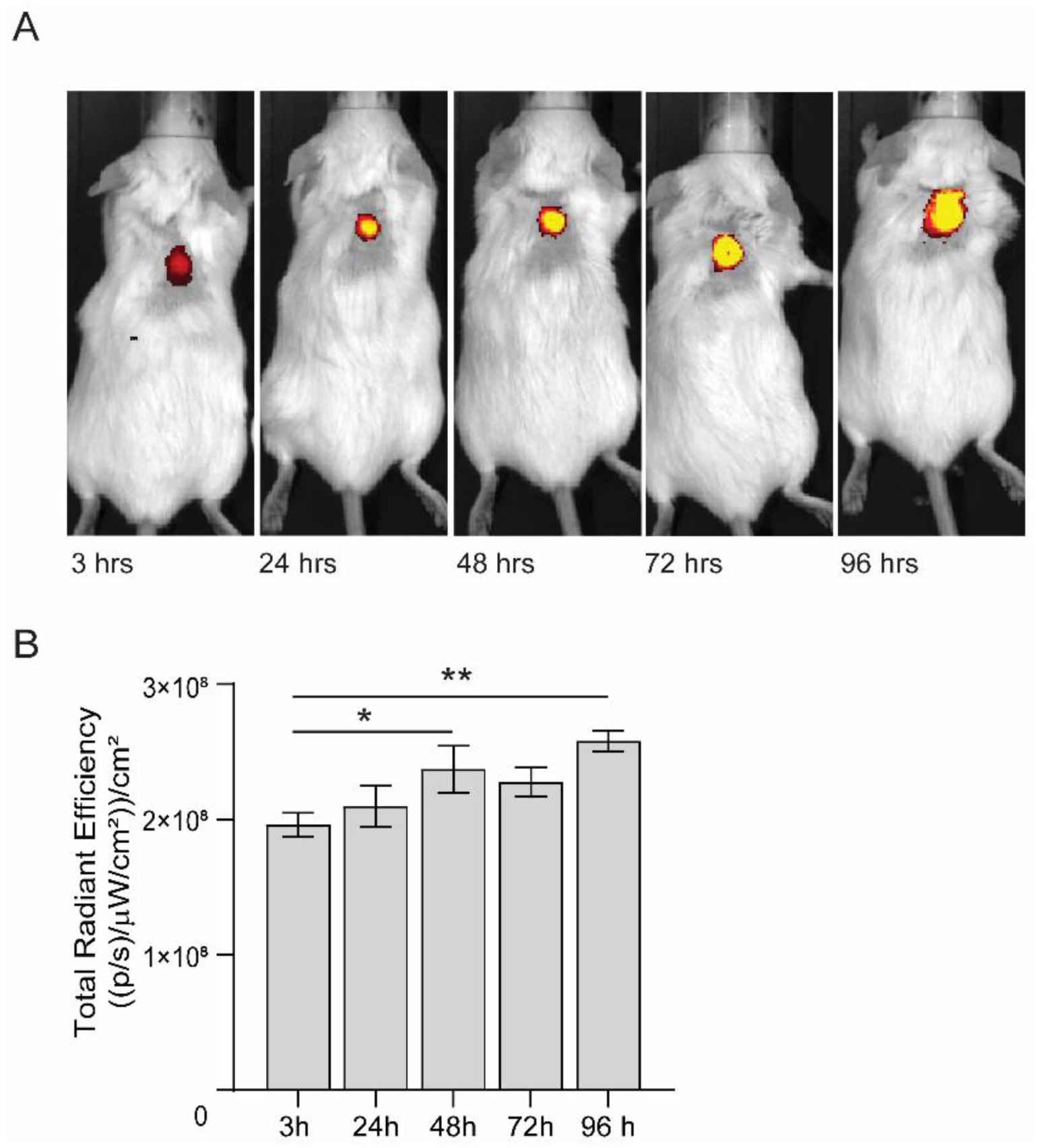
*In vivo* imaging of TTA-UC-PLGA-NPs. (a) FVB mice were inoculated in the back with 1×10e^6^ MCF-7 cells. When the tumors reached a size of ∼125 mm^3^, mice were injected intravenously in the tail vein with 0.5mg of TTA-UC-PLGA-NPs and imaged 3, 24, 48, 72 and 98 hours post injection using the IVIS imaging system. (A) TTA-UC-PLGA-NPs were excited at a wavelength of 535 nm and the emission (phosphorescence) was collected at 640 nm. (B) The total radiance efficiencies were calculated in the tumor areas and the values were compared. Statistical analysis was performed using Mann-Whitney test, * =p 0.026, **= p 0.0022 (6 mice per group were used).

### Measurement of TTA-UC process ex vivo

96 hours post TTA-UC-PLGA-NPs injection, breast adenocarcinoma tumors were excised and the presence of TTA-UC phenomenon was analysed *ex vivo*. To this end, isolated tumors were further processed for cryosectioning and imaged using a confocal microscope. The sections were excited at 514 nm and emissions were collected between 430-475 nm (to observe TTA-UC) and 650-670 nm (to observe phosphorescence) (Figure 9). Strikingly, TTA-UC and phosphorescence signals could be detected in the tumor margins and vessel-like structures 96 hours post injection of TTA-UC-PLGA-NPs. The staining pattern suggests that TTA-UC-PLGA-NPs diffused into the tumor via the margins and entered into deeper tumor areas via leakages in the tumor vasculature. The data further confirmed that, despite initial hydrolysis of the PLGA copolymers (Figure 7), the integrity of TTA-UC-PLGA-NPs was maintained for at least 96 hours *in vivo*.

**Figure 9.**
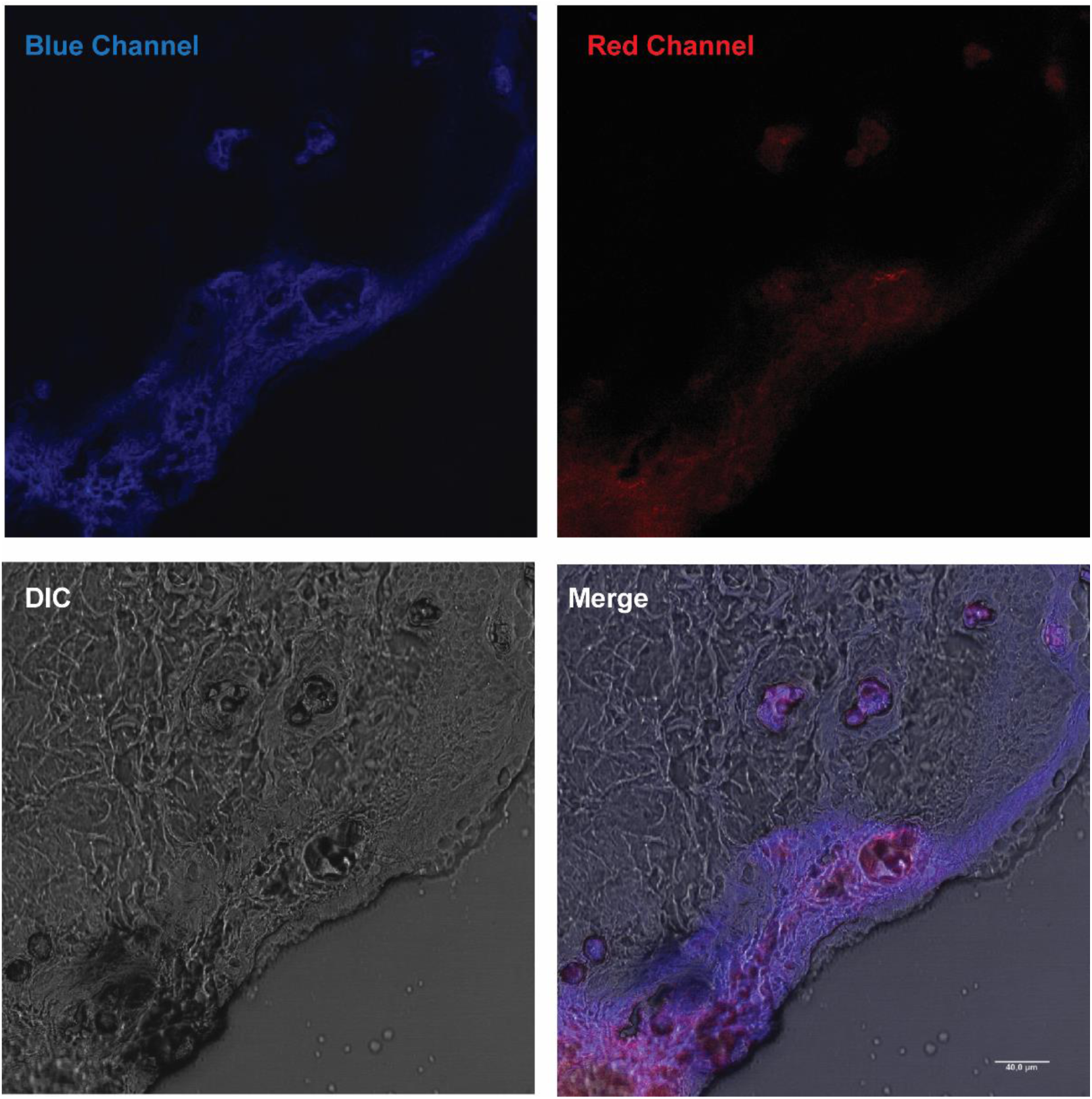
*Ex vivo* imaging of TTA-UC-PLGA-NPs. MCF-tumors were excised 96 hours post injection of TTA-UC-PLGA-NPs and tumor cryosections were imaged by confocal microscopy. All cryosections were excited at 514 nm and the upconversion signal (blue channel) was collected between 430-475 nm and the phosphorescence signal (red channel) between 650-670 nm. Scale bar = 40µm.

In conclusion, in the present study we developed a TTA-UC-PLGA-NP system by incorporating the polycyclic aromatic hydrocarbon DPA and the metalporphyrin PtOEP into a core of hydrophobic PLGA-copolymers. *In vitro* and *ex vivo* studies confirmed the presence of TTA-UC phenomenon resulting from TTA-UC-PLGA-NPs. TTA-UC could be detected 96 hours post injection of TTA-UC-PLGA-NPs in the tumor area *in vivo*, confirming the integrity and suitability of PLGA-NPs as *in vivo* delivery system for TTA-UC, while maintaining the optical integrity of the TTA-UC process. This study provides proof-of-concept evidence on the feasibility of combining the advantageous properties of PLGA for drug delivery with state-of-art TTA-UC for simultaneous optical *in vivo* imaging. The here presented methodology could form the basis for novel TTA-UC systems encapsulated into PLGA, which can be excited in the NIR-I or NIR-II range for clinical translation.

### Experimental Procedures

#### Materials

PtOEP, DPA, DCM and DMF were purchased from Sigma-Aldrich®, The Netherlands. PLGA 50:50 (PLA/PGA) and water-soluble surfactant, polyvinyl alcohol (PVA), were purchased from Evonik industries AG, Germany. MTS (3-(4,5-dimethylthiazol-2-yl)-5-(3-carboxymethoxyphenyl)-2-(4-sulfophenyl)-2H-tetrazolium, inner salt) reagent was purchased from Promega, The Netherlands. Histomount™, agar scientific, United Kingdom. All aqueous solutions were prepared with Milli-Q water.

#### Synthesis of TTA-UC PLGA-NPs loaded with metalloporphyrins and polycyclic aromatic hydrocarbons

The TTA-UC-PLGA-NPs were synthesized according to an oil-in-water emulsion and a solvent evaporation method. Briefly, 100 mg of PLGA and 1 mg of each chromophore were dissolved in 3 ml of DCM. The mixture of chromophores and PLGA polymer was emulsified under sonication (ultrasound tip Branson, Sonifier 250), with 25 ml of water containing 500 mg of PVA for 120 s. The newly formed emulsion was left overnight in stirring at 4°C in order to remove the organic solvent. The PLGA-NPs were collected by centrifugation at 14800 rpm for 20 min, washed three times and freeze-dried. The NPs were stored at 4°C and rehydrated prior to use.

#### Evaluation of physicochemical properties of the TTA-UC-PLGA-NPs

Z-average size, PDI and zeta potential of TTA-UC-PLGA-NPs were measured using a Malvern ZetaSizer 2000 (software: ZetaSizer 7.03). Fixed scattering angle of 90° at 633 nm was set up for the analysis. The measurements were performed after reconstitution of freeze-dried NPs in Milli-Q water.

#### Quantification PtOEP and DPA encapsulation efficiency

The encapsulation efficiency of chromophores was determined by UV/VIS spectrophotometry using an Ultrospec 2100 pro by Amersham Bioscience. Biodegradable PLGA-NPs were hydrolysed overnight with 0,8M NaOH at 37°C. Concentration of dyes was calculated using a calibration curve built with known amounts of PtOEP and DPA.

#### Cell culture

The human ovarian carcinoma cell line OVCAR-3 and the breast cancer cell line MCF**-7** were cultured in RPMI (Roswell Park Memorial Institute 1640 Medium) medium supplemented with 10% of fetal calf serum. The cells were left to attach to the bottom of the plate for 4 hours before treatment with NPs. After 4 hours, the NPs were added to the cells at various concentrations of 25, 50, 100, and 200 µg/ml. DMSO at 25% was used as a positive control. The cells treated with NPs were incubated at 37°C under 5% CO_2_ for 72 hours. After 72 hours the medium was changed and 20 µl of MTS reagent was added to each well. The ELISA reader (Molecular Devices VERSAmax Tunable Microplate Reader, Software: SoftMax Pro v5.4.1) was used to measure the absorbance value of MTS product at 490 nm when the colour of the medium changed from yellow to light brown. The following formula was applied to calculate the viability of cells growth: cell viability (%) = (mean of absorbance value of treated sample/ mean of absorbance of negative (live) control) × 100.

#### In vitro uptake of TTA-UC-PLGA-NPs by OVCAR-3 cells

Fluorescence imaging of fixed OVCAR-3 cells incubated with TTA-UC-PLGA-NPs was performed with a Leica DMRA microscope using a PL APO 63×/1.32-0.6 oil objective. DAPI and Cy5 filters were used for nucleus and membrane detection, respectively, using the following protocol: Adherent cells were detached with 0.2% trypsin in PBS (Gibco by Life Technologies, The Netherlands) and seeded in 8-well chamber slides (BD Biosciences, The Netherlands) at a concentration of 2×10^4^ cells/well. After 4 hours, the TTA-UC-PLGA-NPs were added at a concentration of 200 μg/ml for 2 hours at 37°C under 5% of CO_2_. At the end of the incubation time, the cells were washed with PBS, fixed in 4% of paraformaldehyde (PFA) and stained with DiD labelling solution according to manufacturer protocol (Invitrogen, The Netherlands). The staining was completed adding Vectashield mounting medium containing DAPI.

#### Measurement of UC process

Confocal microscopy imaging of TTA-UC-PLGA-NPs was performed by Leica SP8 X WLL (White Light Laser) laser scanning microscope. To this end, TTA-UC-PLGA-NPs were dissolved at 5 µg/µl in distilled water and dropped onto a glass slide and imaged. Emissions were collected between 650-670 nm and 430-475 nm; the excitation was provided at 535nm.

#### Stability measurement of TTA-UC-PLGA-NPs in solution

In order to evaluate the stability of the TTA-UC system inside the PLGA-core, the TTA-UC-PLGA-NPs were dissolved in PBS and left for 0, 24 and 72 hours shaking (300 rpm) at 37°C. At each time point phosphorescence and UC-luminescence were measured using a spectrofluorometer (Fluorolog). The suspension of TTA-UC-PLGA-NPs was excited at 535 nm and the emission signals were captured at 433 and 641 nm.

#### Animals

Female FVB mice were purchased from Charles River Laboratories (L’Arbresle Cedex, France). Animals were housed at 22°C and 50% of humidity with free access to food and water, and maintained under standard 12 hours light/12 hours dark cycles.

All animal experiments were assessed according to the ethics of animal research and approved by the Animal Welfare Committee of Leiden University Medical Center, the Netherlands. All mice received humane care and were kept in compliance with the Code of Practice Use of Laboratory Animals in Cancer Research (Inspectie W&V, July 1999). All analytical procedures were performed under isoflurane gas anaesthesia (3% induction, 1.5-2% maintenance) in 70% pressurized air and 30% O_2_, unless stated differently.

#### In vivo monitoring of TTA-UC-PLGA-NPs

*In vivo* imaging was performed using the IVIS spectrum Preclinical Imaging System (Caliper LS, Hopkinton). The images were analysed with Living Image 4.3.0 software.

First, the tumor was induced by injecting 1×10^6^ MCF cells in 100 µl PBS subcutaneously into the back of the mice. When the tumors reached the volume of approximately 125 mm^3^, 0.5 mg TTA-UC-PLGA-NPs in 100 µl PBS were injected into the tail vein of each mouse. At 3, 24, 72 and 96 hours post injection optical imaging of treated mice was performed. Biodistribution kinetics at each time point were measured by quantifying the fluorescence intensity in pre-set regions of interest (ROI) at the tumor site expressed as the average radiant efficiency in (p/sec/cm^2^/sr/)/(µW/cm^2^).

#### Measurement of TTA-UC ex vivo

After 96 hours, the mice were sacrificed and subcutaneous tumors were surgically removed. Freshly isolated tumors were placed in tissue moulds and covered with Tissue-Tek^®^ O.C.T.™ (Sakura). For snap freezing, the tumors were left on dry ice for some minutes and then stored at −80°C. Cryosectioning of tumors was performed using CryoStar™ NX70 at the working temperature between −25 and −30°C, the thickness of the sections was fixed between 5 and 14 µm. Before microscopic analysis of the tumors, the sections were washed in Milli-Q water, dried under the hood and mounted in medium containing Mowiol (Sigma) and 2.5% DABCO (Sigma).

#### Statistical data analysis

Graph Pad Prism software version 7 was used to perform statistical analysis. Mann-Whitney test was applied in all experiments.

## Acknowledgements

O.V. and L.J.C. were supported by project grants from the European Commission H2020-MSCA-RISE (644373 - PRISAR), H2020-MSCA-RISE (777682 - CANCER), H2020-WIDESPREAD-05-2017-Twinning (807281 - ACORN), H2020-WIDESPREAD-2018-03 (852985 - SIMICA), H2020-SCA-RISE-2016 (734684 - CHARMED) and MSCA-ITN-2015-ETN (675743-ISPIC), 861190 (PAVE), 857894 (CAST), 859908 (NOVA-MRI); 860173 (RISE-WELL); 872860 (PRISAR2). L.J.C was also supported by the research program VIDI (project number 723.012.110) of Dutch Research Council (NWO). C.E. was supported by the research program VENI with project number 916.181.54, which is (partly) financed by the Dutch Research Council (NWO) and the European Union’s Horizon 2020 research and innovation programme under the Marie Sklodowska Curie grant agreement SIMICA (852985).

## Abbreviations

DCM: Dichloromethane;
DLS: dynamic light scattering;
DMF: dimethylformamide;
DPA: 9,10-diphenylanthracene;
EE: encapsulation efficiency;
ISC: intersystem crossing;
NPs: nanoparticles;
PDI: polydispersity index;
PLGA: poly(lactic-co-glycolic acid;
PtOEP: Platinum(II) octaethylporphyrin;
TTA: triplet-triplet annihilation;
TTET: triplet-triplet energy transfer;
UC: upconversion.

## Notes

### Competing Interest Statement

The authors have declared no competing interest.

